# A comparative ultrastructure study of the tardigrade *Ramazzottius varieornatus* in the hydrated state, after desiccation and during the process of rehydration

**DOI:** 10.1101/2023.05.03.539216

**Authors:** Simon Galas, Emilie Le Goff, Chantal Cazevieille, Akihiro Tanaka, Pierre Cuq, Stephen Baghdiguian, Takekazu Kunieda, Nelly Godefroy, Myriam Richaud

## Abstract

Tardigrades can survive hostile environments such as desiccation by adopting a state of anhydrobiosis. Numerous tardigrade species have been described thus far, and recent genome and transcriptome analyses revealed that several distinct strategies were employed to cope with harsh environments depending on the evolutionary lineages. Detailed analyses at the cellular and subcellular levels are essential to complete these data. In this work, we analyzed a tardigrade species that can withstand rapid dehydration, *Ramazzottius varieornatus*. Surprisingly, we noted an absence of the anhydrobiotic-specific extracellular structure previously described for the *Hypsibius exemplaris* species. Both *Ramazzottius varieornatus* and *Hypsibius exemplaris* belong to the same evolutionary class of Eutardigrada. Nevertheless, our observations reveal discrepancies in the anhydrobiosis mechanisms between these two species. Interestingly, these discrepancies are correlated with their variations in dehydration resistance.

## 1. Introduction

Tardigrades are tiny metazoan animals that range in size from approximately 0.1–1.2 mm and have four pairs of legs. They can be called “water bears” because of their appearance and “moss piglets” because of where they can be found. Nearly 1338 tardigrades species have thus far been described [1], which are distributed from the depths of the oceans to the highest mountain peaks [2]. The worldwide distribution of tardigrade species can be either endemic or cosmopolitan [3], and their transport by birds or snails has recently been suggested [4–6].

As earlier as their discovery in the 18th century [7], tardigrades have demonstrated an ability to adopt a latent state due to a shortage of water, which is called anhydrobiosis. Tardigrades can enter an anhydrobiosis state in response to desiccation to form an anhydrobiote, allowing the organism to wait for the return of water. Thus, by reaching nearly complete desiccation, tardigrades can survive for up to 10 years as anhydrobiotes [8,9]. During the course of desiccation, tardigrades contract and retract their whole body to assume a characteristic “tun”-shaped anhydrobiote structure. Tardigrades can then lose up to 97% of their bound and free body water content. Upon reaching the anhydrobiotic state, tardigrades enter a cryptobiosis phase that is characterized by an ametabolic status [10].

Other invertebrates, such as rotifers, nematodes and dipteran larvae [11–13], can also enter anhydrobiosis. However, tardigrades are distinguished from other invertebrates by their resilience to extreme stresses, such as temperatures ranging from -272 to +150°C [14,15], very high pressures (up to 7.5 GPa) equivalent to that at a depth of up to 180 km from the Earth’s surface [16], radiation at levels up to 5000 Gy [17–19] and exposure to solar radiation at a low Earth orbit in a space vacuum during a ten-day space flight [20].

To date, the genomes of four tardigrade species are available [21]. The genomes of two Eutardigrada species: *Ramazzottius varieornatus* and *Hypsibius exemplaris* [22–24], enabled the identification of gene products involved in anhydrobiosis. For instance, the Dsup (damage suppressor) gene was identified in *Ramazzottius varieornatus* (*R. varieornatus*) and was suggested to protect both human and plant cells from gamma ray irradiation [18,24,25] as well as human cultured cells from oxidation by free radicals [18,24]. The molecular capacity of the Dsup gene products to protect nucleosomes from direct oxidation by hydroxyl radicals was thereafter evidenced by an in vitro assay [26].

While *R. varieornatus* is tolerant to a rapid desiccation process (minutes), *H. exemplaris* can undergo effective anhydrobiosis after only an obligate preconditioning period (hours) [21,27]. In accordance with, both species show contrasting gene expression in response to desiccation [21,28]. Thus, while *R. varieornatus* is believed to express anhydrobiosis involved genes constitutively, *H. exemplaris* species requires a *de novo* expression induction of orthologs genes [21]. To date, tardigrade species belonging to the Heterotardigrada class seem to lack bona fide Dsup orthologs [26].

It has been shown [29] that tardigrades belonging to the Eutardigrada class, such as *R. varieornatus* and *H. exemplaris*, also possess genes [24,30,31] encoding proteins that are involved in resistance to desiccation stress, such as the cytosolic abundant heat-soluble (CAHS), secretory abundant heat-soluble (SAHS), late embryogenesis abundant mitochondrial (RvLEAM), and mitochondrial abundant heat-soluble (MAHS) proteins. These intrinsically disordered proteins (IDPs) are involved in the maintenance of cellular structures during desiccation processes [24,30,32–34].

Species of the Eutardigrada class possess specific genes involved in stress resistance that differ from those of other Tardigrada classes, such Heterotardigrada, but some discrepancies have also been reported among Eutardigrade species. For example, *R. varieornatus* possesses a trehalose-6-phosphate synthase gene, while *H. exemplaris* does not [28]. Trehalose-6-phosphate synthase can produce the nonreducing sugar trehalose [35], which has been proposed to play a role in mediating desiccation tolerance in some organisms, such as *Caenorhabditis elegans, Saccharomyces cerevisiae* and chironomids, by vitrifying their cellular contents. However, other desiccation-tolerant invertebrates, such as rotifers, do not require this sugar [36–39] and the presence of the trehalose is still unclear in tardigrades [40–44].

To date, only a few ancient reports have attempted to describe the ultrastructures of anhydrobiotic organisms [45–47]. Several analyses of hydrated tardigrades have been conducted [48–60]. However, studies on the ultrastructure of tardigrades in the anhydrobiotic tuns are extremely scarce. Halberg *et al*. [61] described the tun morphology of the *Richtersius coronifer* species with an emphasis on muscular organization, while Czernekova *et al*. [62,63] investigated the internal morphologies of dehydrated organs, tissues and cells in the same species.

In a previous report [64], we used electron microscopy to compare hydrated and anhydrobiotic tuns of *H. exemplaris*. We highlighted deep modifications occurring up to the subcellular level in the anhydrobiote and during the course of exit from anhydrobiosis. We also uncovered the materialization of an anhydrobiote-specific and reversible extracellular structure [64].

In the present study, we studied the structures and ultrastructures of the cells and organelles of anhydrobiotic *R. varieornatus* specimens by electron microscopy and compared them to the ultrastructures of active hydrated specimens. Finally, we compare strategies used by the Eutardigrada species *R. varieornatus* and *H. exemplaris* to resist anhydrobiosis.

## 2. Materials and Methods

### Materials

The Yokozuna-1 strain of the extremotolerant *R. varieornatus* Bertolani and Kinchin [65], (Eutardigrada, Hypsibiidae), provided by Takekazu Kunieda (University of Tokyo), was used for all experiments. Tardigrades were cultured as previously described [14]. They were fed with the unicellular algae *Chlorella vulgaris (*strain A60) on 2% Bacto agar plates prepared with Volvic water and incubated at 20°C under constant dark conditions. Algae were purchased from the Sciento Company (Manchester, UK).

### Desiccation protocol

Twenty specimens in a drop of water were placed on a filter paper inside Petri dish, which were left at room temperature (20-22°C) and relative humidity (RH) (between 30-36%) for one hour to confirm good dehydration. The dessication process of the specimens was monitored by direct observation under a stereomicroscope in order to assure that the tardigrades underwent a proper anhydrobiosis process and formed a tun. The anhydrobiotes were stored at 20°C and at room RH (between 30-36%) in an incubator for one week before analysis.

### Rehydration protocol

To rehydrate the desiccated *R. varieornatus* after one week of dehydration, Volvic water droplets were added to the filters. Tardigrades were maintained in water at room temperature (20-22°C) and prepared for TEM after 5 and 15 minutes of contact with liquid.

### Scanning electron microscopy

Tardigrades were fixed with 2.5% glutaraldehyde in PHEM buffer (pH 7.2) for one hour at room temperature and then washed with PHEM buffer. The fixed samples were dehydrated using a graded ethanol series (30-100%), followed by 10 minutes incubations in graded ethanol-hexamethyldeliazane and then hexamethyldeliazane alone. Subsequently, the samples were sputter coated with a gold film that was ap-proximately 10 nm thick and then examined under a scanning electron microscope (Hitachi S4000, at CoMET, Institut des Neurosciences Montpellier, MRI-RIO Imaging, Biocampus Montpellier France) using a lens detector with an acceleration voltage of 10 kV at calibrated magnifications.

### Transmission electron microscopy

According to Richaud et al. [64], samples were fixed in 2.5% glutaraldehyde in PHEM buffer (1X, pH 7.4) overnight at 4°C, rinsed in PHEM buffer and postfixed in 0.5% os-mic acid for 2 hours in the dark at room temperature. After two rinses in PHEM buffer, the samples were dehydrated in a graded series of ethanol (30-100%) and embedded in EmBed 812 using an automated microwave tissue processor for electronic microscopy (Leica EM AMW). Ultrathin sections (70 nm; Leica-Reichert Ultracut E) were collected from different levels of each block, counterstained with 1.5% uranyl acetate in 70% ethanol and lead citrate and observed using a Tecnai F20 transmission electron micro-scope at 200 kV at the CoMET MRI facilities (INM, Montpellier, France). For TEM, 5 tardigrades were analyzed for each condition: tuns, rehydrated for 5 minutes, rehy-drated for 15 minutes and hydrated.

### Tardigrade size

We measured the sizes of hydrated and tun tardigrades from the tip of the head to the extreme end of the body. Measurements from DIC images obtained using a Zeiss LSM880 Fast Airyscan confocal microscope at the DBS-Optique MRI facilities (Montpellier, France) were determined with ImageJ software. For each condition, five speci-mens were measured.

### Number of nuclei

We counted the number of nuclei in hydrated and tuned animals. Counts from DAPI z-stack stained images obtained using a Zeiss LSM880 Fast Airyscan confocal micro-scope at the DBS-Optique MRI facilities (Montpellier, France) were determined with ImageJ software. For each condition, the number of nuclei was counted in five tardigrades.

### Mitochondrial size

We measured the mitochondrial sizes under every condition: hydrated, dehydrated and after 5 or 15 minutes of rehydration. For each condition, mitochondria were observed in each cell types and in the same proportions to avoid sampling bias. The sizes of 140, 150, 110 and 136 mitochondria were measured in each group, respectively, using ImageJ software. Mitochondria were measured in cross-sections through and through.

### Statistical analysis

According to Richaud et al. [64], we used XLSTAT software (Addinsoft, New York, NY, USA) to compare mitochondrial sizes among animals that were hydrated, rehydrated for 5 or 15 minutes and dehydrated and to compare the body sizes and numbers of nuclei between hydrated and dehydrated animals.

## 3. Results

### 3.1. Comparison of hydrated and anhydrobiotic Ramazzottius varieornatus

#### 3.1.1. Cell compaction of anhydrobiotic tardigrades

To compare the ultrastructures and cell shapes of hydrated and anhydrobiotic tardigrades, groups of *R. varieornatus* were dehydrated for one week and then analyzed for comparison with control active hydrated groups. We first used scanning electron microscopy (SEM) to obtain a global view of the external morphologies of hydrated versus anhydrobiotic tuns. Figure 1a and b shows representative images of the characteristic contraction in anhydrobiotic tun specimens.

**Figure 1.**
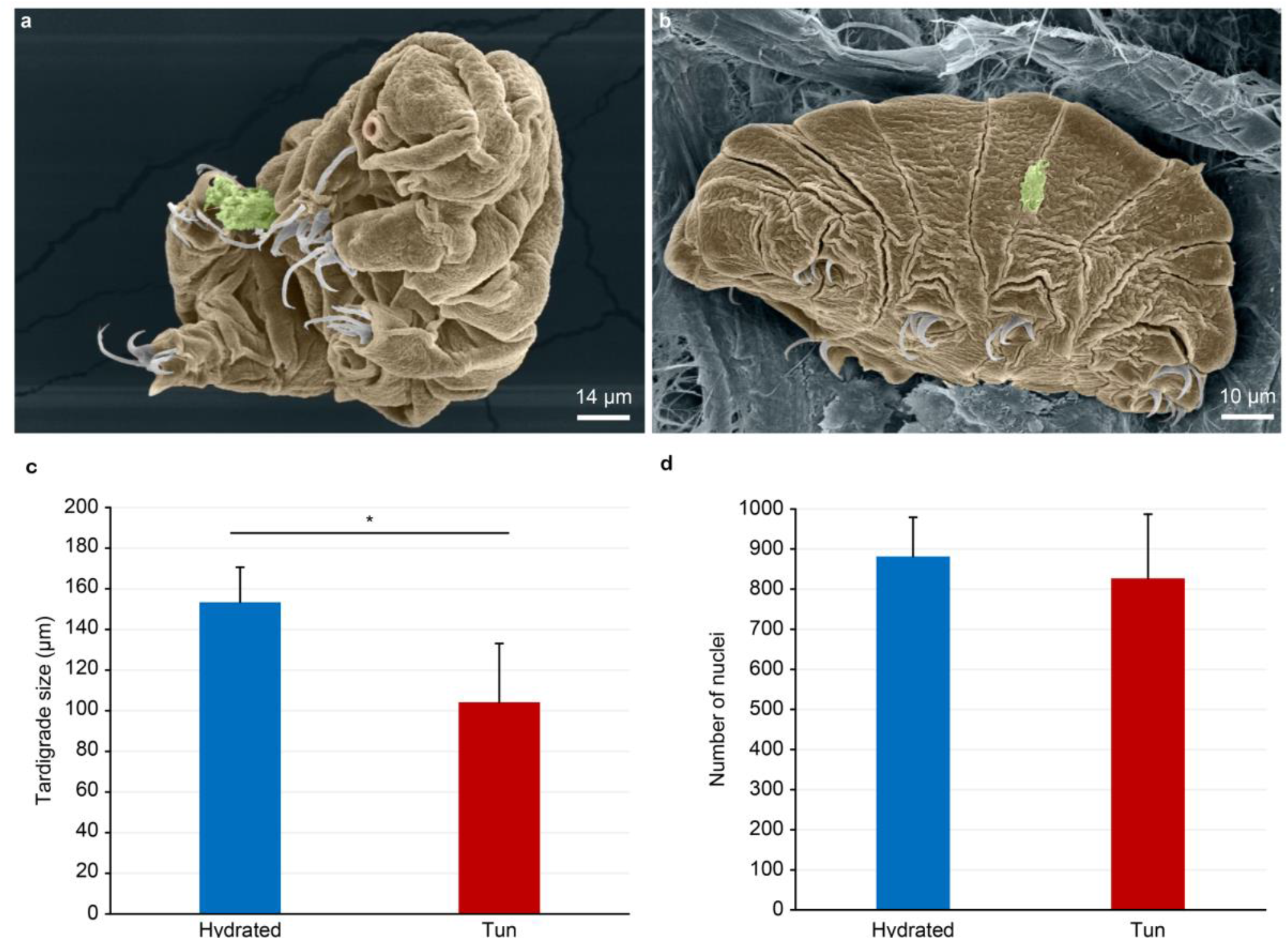
Comparison of hydrated and anhydrobiotic *R. varieornatus:* (**a**–**b**) SEM images. (**c**–**d**) Statistic analyses of body sizes and nucleus numbers. (**c**) The error bars indicate the standard deviation, and the stars indicate significant difference (Kolmogorov-Smirnov test, p = 0.048; α = 0.05). (**d**) The error bars indicate the standard deviation.

To access targeted internal structures with nuclear-specific dyes, we performed confocal laser microscopy and made observations with differential interference contrast (DIC). Length measurements of hydrated specimens revealed an average size of 153 +/- 17 μm, while the anhydrobiotic tuns showed an average size of 104 +/- 29 μm (Figure 1c), revealing a size reduction of 32% (Figure 1c). However, staining the nuclei of both the hydrated and anhydrobiotic groups with DAPI revealed a nonsignificant difference in the total cell counts (Figure 1d).

#### 3.1.2. Comparative analysis of cell structure and ultrastructure

To better understand the origin of anhydrobiotic tun stress resistance, we performed transmission electron microscopy (TEM) analyses of tissues from five individuals in each condition.

The hydrated individuals showed a large space between cells, named extracellular space (es) (Figure 2a). The sizes of this space throughout the body of the tardigrade were not similar. Epidermal cells, bordered by the cuticle, were clearly visible together with numerous pigmented vesicles (Figure 2c).

**Figure 2.**
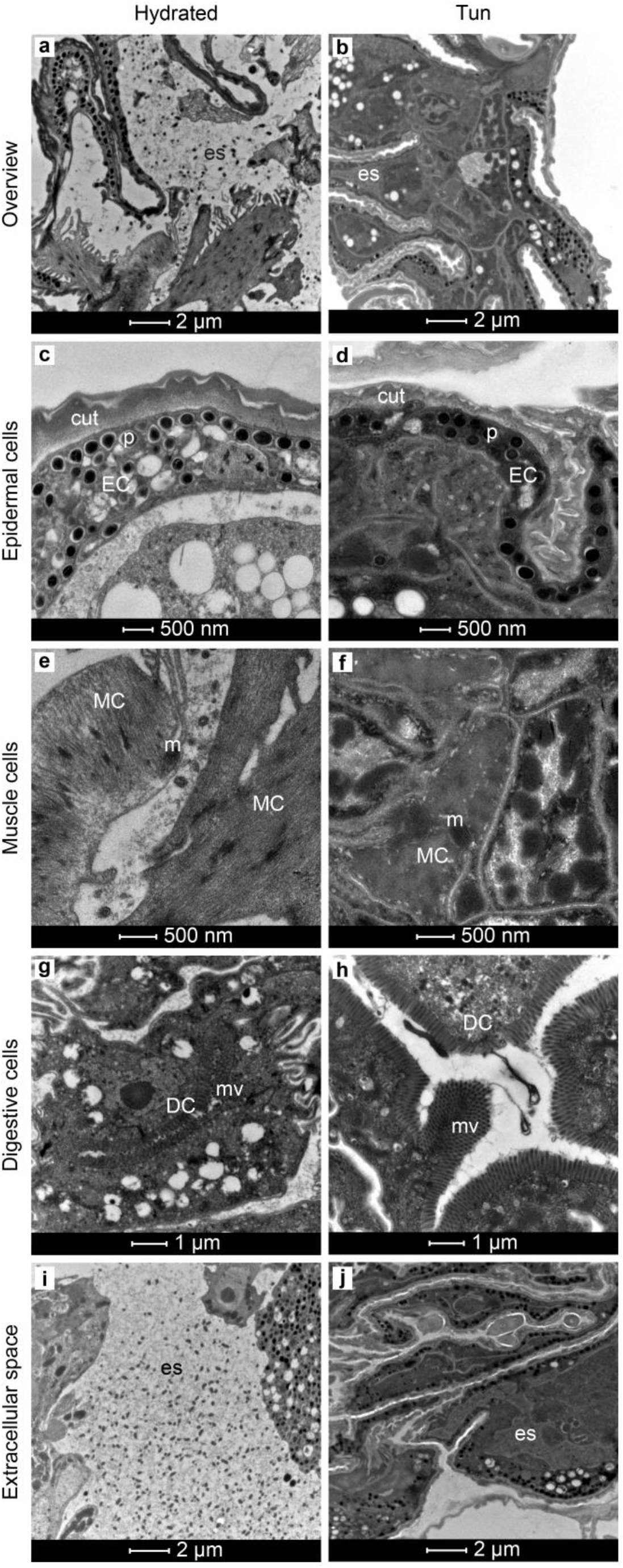
Ultrastructures of *R. varieornatus* under the hydrated and tun statuses. (**a, b**) Overview of a body part. (**c, d**) Ultrastructure of epidermal cells. (**e, f**) Ultrastructure of muscle cells. (**g, h**) Ultrastructure of digestive cells. (**i, j**) Extracellular space overview. Cut: cuticle, DC: digestive cell, EC: epidermal cell, MC: muscle cell, m: mitochondria, mv: microvilli, p: pigment, es: extracellular space.

The muscle cells possessed long fibers with long dark mitochondria (Figure 2e), and digestive cells appeared with long villosities (Figure 2g). Additionally, numerous lipid droplets were observed inside the cells regardless of the cell type.

While we also observed es (Figure 2b) features in the anhydrobiotic tun group, they sometimes appeared narrower than those in hydrated individuals, probably due to tardigrade compaction (Figure 2b). We observed a space between the epidermal cells and the cuticle (Figure 2d), but the global structure of this cell type was not affected by dehydration (Figure 2d). Conversely, muscular fibers were less discernible in the anhydrobiotic tun group than in the hydrated group because of compaction (Figure 2f). Similar to the hydrated tardigrades, the digestive cells of the anhydrobiotic group exhibited long villosities (Figure 2h), and lipid droplets were still present. Surprisingly, no apoptotic cells were observed in any of the specimens.

In both the hydrated and anhydrobiotic tun groups, dot-like structures were observable in the es (Figure 2i and j), which appeared to be more condensed in anhydrobiotic tardigrades than in hydrated groups; however, their number seemed roughly comparable.

The hydrated and anhydrobiotic groups showed comparable organelle structures, with numerous mitochondria being observed in both groups. Moreover, the mitochondrial structures were comparable (Figure 3a and c vs b and d) and not degraded. Surprisingly, the mitochondrial cristae in the anhydrobiotic tun group were comparable to those in hydrated animals (Figure 3c and d); however, a statistically significant size reduction of 24% (Figure 3e) was observed in mitochondria of the anhydrobiotic tun group compared with the hydrated group.

**Figure 3.**
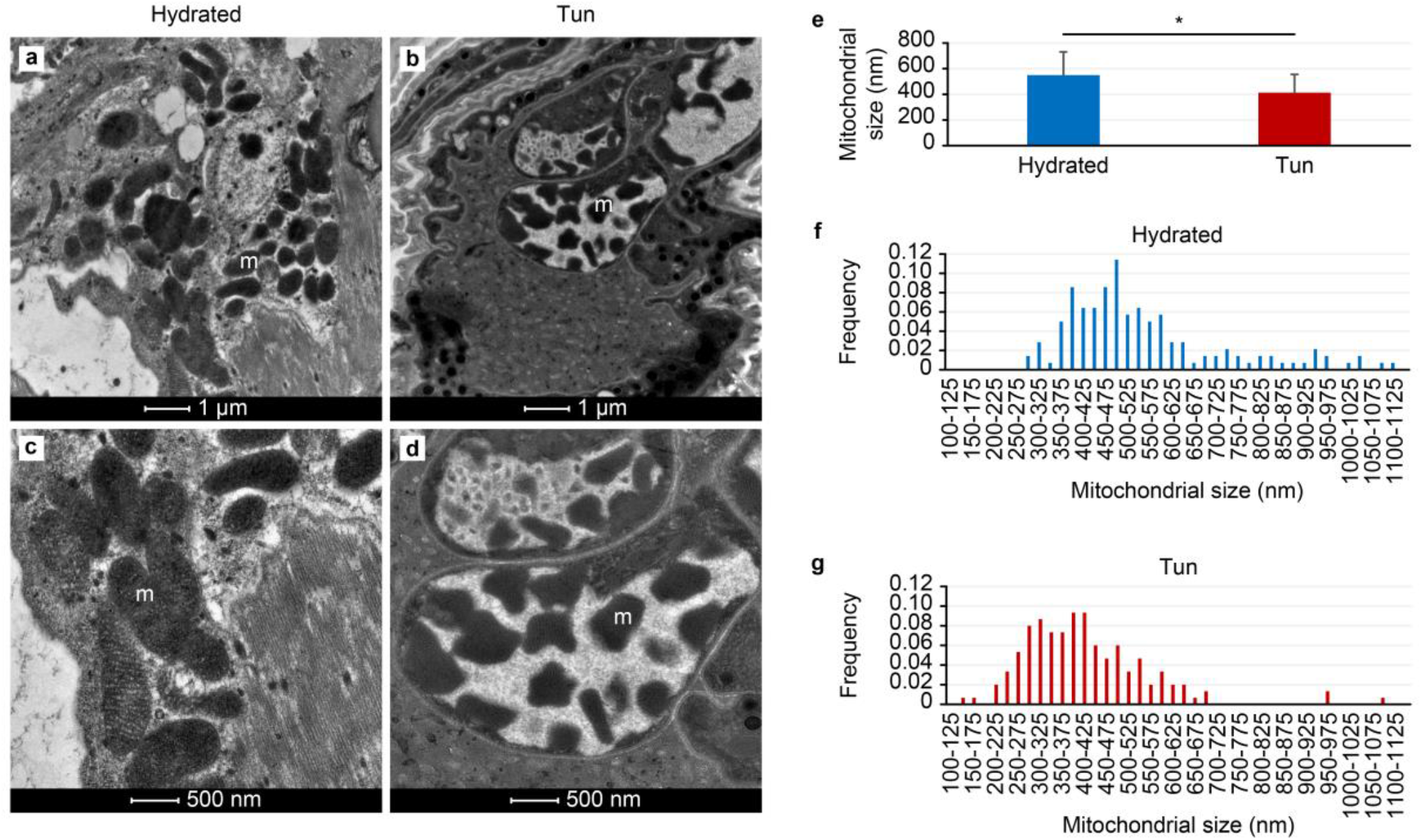
Comparison of mitochondria in hydrated and anhydrobiotic *R. varieornatus* cells. (**a**-**d**) Transmission electron microscopy images. (**e**) Mean mitochondrial size. The error bars indicate the standard deviation, and the stars indicate significant differences (Student’s t-test, α = 0.05). Table 1 shows the complete statistical results. (**f, g**) Histogram of the mitochondrial size frequencies of hydrated tardigrades (**f**) and desiccated tardigrades (**g**). m: mitochondria.

Figure 3f and g shows the size distribution frequencies of up to 140 and 150 mitochondria measured in both the hydrated and anhydrobiotic tun groups respectively. Additionally, we observed many mitochondria with atypical shapes in the anhydrobiotic tun group (Figure 3b and d) that were not observable in the hydrated group (Figure 3a and c).

**Table 1.** Statistical results of the mitochondrial sizes (nm) in cells from *R. varieornatus* in 4 stages: tuns (A), after 5 minutes of rehydration (B), after 15 minutes of rehydration (C) and hydrated (D).

| Stage | Mean ± SD | Code | p-value* | n |
| --- | --- | --- | --- | --- |
| Tun | 413 ± 142 | A | 0.0158 (B)<br>< 0.0001 (C)<br>< 0.0001 (D) | 150 |
| Rehydrated for 5 min | 453 ± 115 | B | 0.0158 (A)<br>< 0.0001 (C)<br>< 0.0001 (D) | 110 |
| Rehydrated for 15 min | 572 ± 136 | C | < 0.0001 (A)<br>< 0.0001 (B)<br>0.2562 (D) | 136 |
| Hydrated | 550 ± 181 | D | < 0.0001 (A)<br>< 0.0001 (B)<br>0.2562 (C) | 140 |

\*Student's t-test, $\alpha=0.05$

### 3.2. Temporal change in anhydrobiotic tuns during rehydration

To better understand the functional structures of stress-resistant anhydrobiotic tuns, we assessed the ultrastructural changes in the anhydrobiotic tun group over the course of rehydration. Because anhydrobiotic tuns take only a few minutes (10-20 minutes) to wakeup (size recovery and detectable movements) from dehydration, they were dehydrated for one week, rehydrated for 5 and 15 minutes and then assessed by TEM.

#### 3.2.1. Rehydration of anhydrobiotic tuns for 5 minutes

After 5 minutes of rehydration in the anhydrobiotic tun group, we observed a size evolution that fell between that of the hydrated and anhydrobiotic specimens. Following this observation, we again noticed persistence of the anhydrobiotic state with decoupling between the epidermal cells and the cuticle (Figure 4a). Moreover, the global structure of the epidermal cells containing the already described vesicles was maintained (Figure 4a versus Figure 2c and d), and muscle cells exhibited a normal structure with long contractile fibers (Figure 4c). Gut cells also showed a normal ultrastructure compared to those of the hydrated group (Figure 4e versus Figure 2g and h). The mitochondrial size was intermediate between those in the anhydrobiotic and hydrated groups of tardigrades (Figure 5b and e). The mitochondrial size differences between the anhydrobiotic and hydrated groups were evident based on their size frequency distributions (Supplementary Data S1). More than 110 mitochondria were assessed, and their intermediate sizes were also confirmed by statistical analysis (Table 1). We observed a higher concentration of mitochondria around muscle fibers. Previously described lipid droplets were still observable inside the cells (data not shown) as were the dot-like structures in the es (Figure 4g).

**Figure 4.**
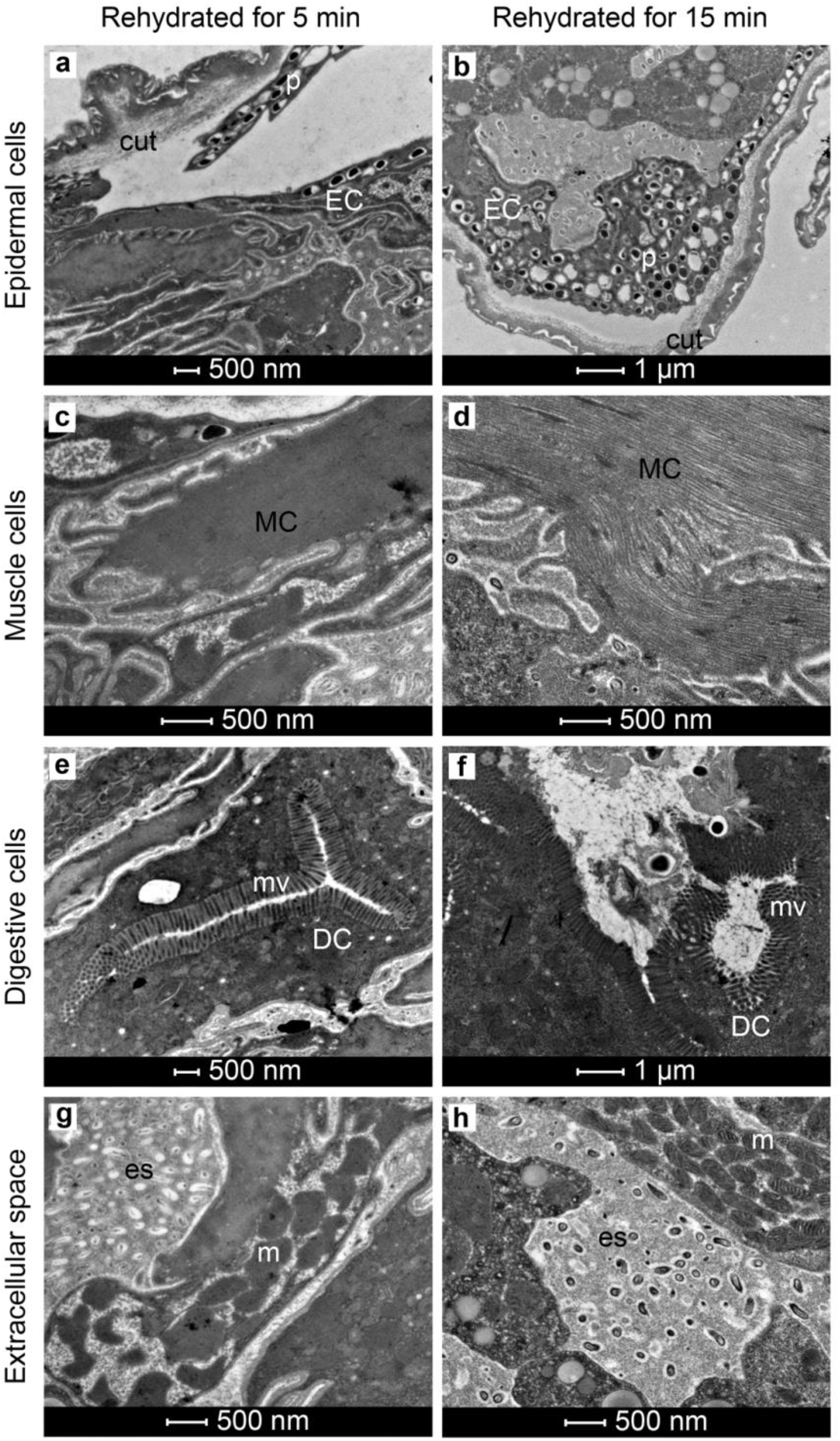
Ultrastructure of *R. varieornatus* after 5 and 15 minutes of rehydration. (**a, b**) Ultrastructure of epidermal cells. (**c, d**) Ultrastructure of muscle cells. (**e, f**) Ultrastructure of digestive cells. (**g, h**) Extracellular space overview. cut: cuticle, DC: digestive cell, EC: epidermal cell, MC: muscle cell, m: mitochondria, mv: microvilli, p: pigment, es: extracellular space.

**Figure 5.**
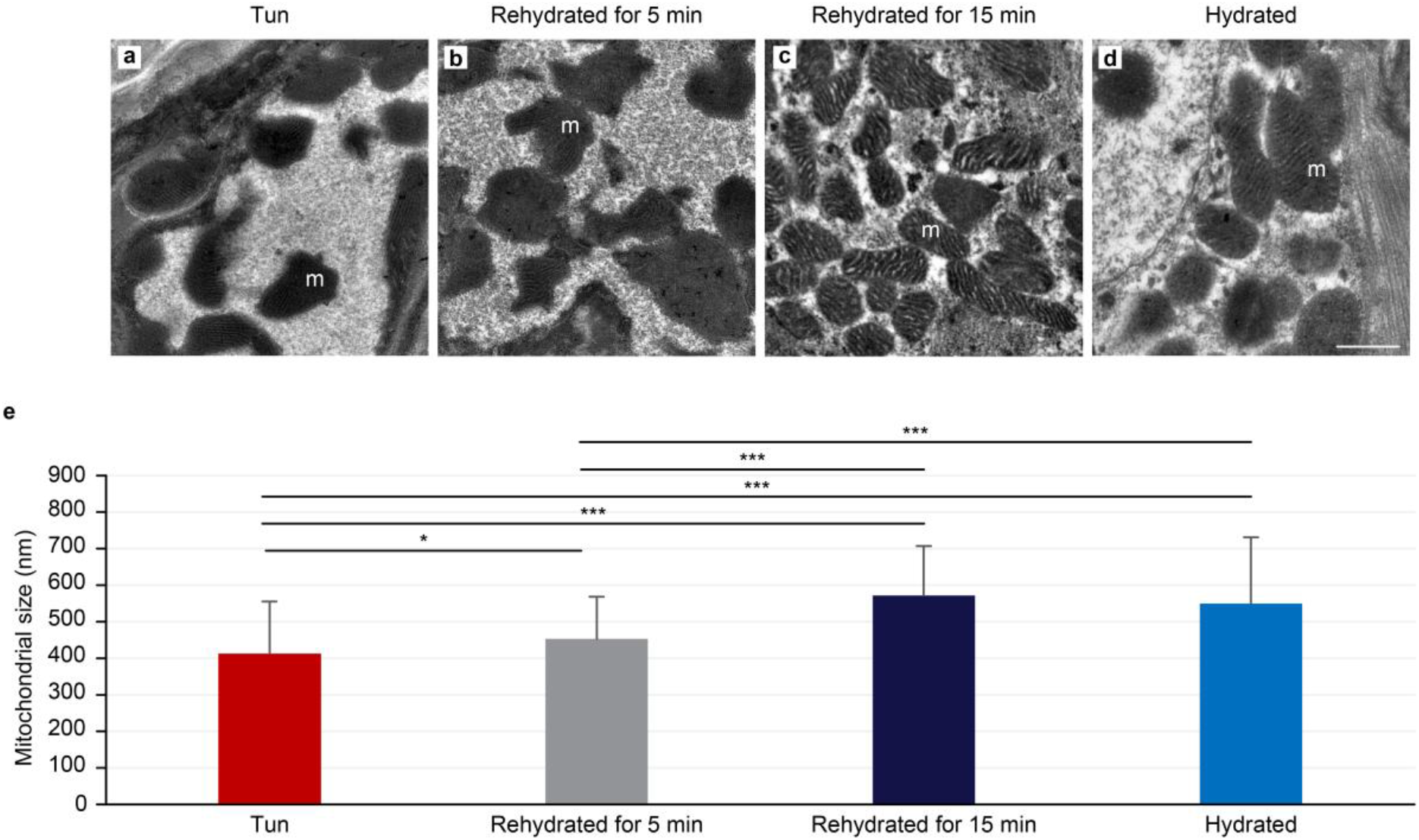
Comparison of mitochondria from *R. varieornatus* in 4 stages: tuns, after 5 minutes of rehydration, after 15 minutes of rehydration and hydrated. (**a**–**d**) Transmission electron microscopy images. Scale bar a-d = 500 nm. (**e**) Mean mitochondrial sizes. The error bars indicate the standard error of the mean. <*> indicates a significant difference at p < 0.05 (Student’s t-test, α = 0.05). <***> indicates a significant difference at p < 0.0001 (Student’s t-test, α = 0.05). See Table 1 for all of the statistical results. m: mitochondria.

#### 3.2.2. Rehydration of anhydrobiotic tuns for 15 minutes

Fifteen minutes after rehydration in the anhydrobiotic tun group, the tardigrade size was already comparable to that of the hydrated group. In agreement with this observation, the epidermal cells recovered contiguously with the cuticle (Figure 4b), and the muscle cells appeared as classical long fibers, like in the hydrated group (Figure 4d versus Figure 4e, 2e and f). In addition, the digestive cells showed a normal structure compared with that in the hydrated group (Figure 4f versus Figure 4e, 2g and h). Moreover, the mitochondrial sizes were equivalent to those in the hydrated control group (Figure 5c and e). This was shown by evaluating the size frequency distribution of up to 136 mitochondria (Supplementary Data S1) and confirmed by statistical analysis (Table 1). Finally, the lipid droplets, previously described in other conditions were still present, as were the dot-like structures in the es (Figure 4h).

## 4. Discussion

*R. varieornatus* can cope with rapid dehydration and is known to be one of the most resilient to desiccation among the limno-terrestrial tardigrades [21,24]. However, no information on internal reorganization during anhydrobiosis is available.

We have previously reported [64] that upon desiccation, *H. exemplaris* shows active secretory cells that are closely related to a specific and reversible extracellular structure surrounding each cell. This specific extracellular structure and the accompanying secretory cells disappear during rehydration, implying their direct association with resistance to dehydration stress. However, *H. exemplaris* is more sensitive to desiccation than *R. varieornatus* [24,28,29] and thus requires preconditioning steps to achieve successful anhydrobiosis [27,32,66].

To determine whether the cell ultrastructural modifications described for *H. exemplaris* [64] are specific or relevant to other tardigrades of the Eutardigrada class, we explored the entry and exit of anhydrobiosis in *R. varieornatus* by electron microscopy.

Recent genomic and transcriptomic comparative analyses [24,28,29,32,35,67] have emphasized genetic differences between *R. varieornatus* and *H. exemplaris*, which include molecular variations in genes that are related to DNA repair [29,67] and anhydrobiosis [28,29]. Moreover, the constitutive expression of genes involved in dehydration-rehydration processes has been proposed to explain the higher resistance of *R. varieornatus* than *H. exemplaris* when facing desiccation [24]. In accordance, during the dehydration process, a major transcriptional commitment was observed in *H. exemplaris*, while limited transcriptional regulation was noticed in *R. varieornatus* [28].

Based on the reported genetic discrepancies and differences on the observed desiccation resistance between the *R. varieornatus* and *H. exemplaris* species, we compared their internal ultrastructural rearrangements during anhydrobiosis. Figure 6 summarizes their ultrastructural divergences during the dehydration process and anhydrobiote formation. Contrary to *H. exemplaris* [64], no secretory cells with a dense network of endocytoplasmic reticulum were found in *R. varieornatus*.

**Figure 6.**
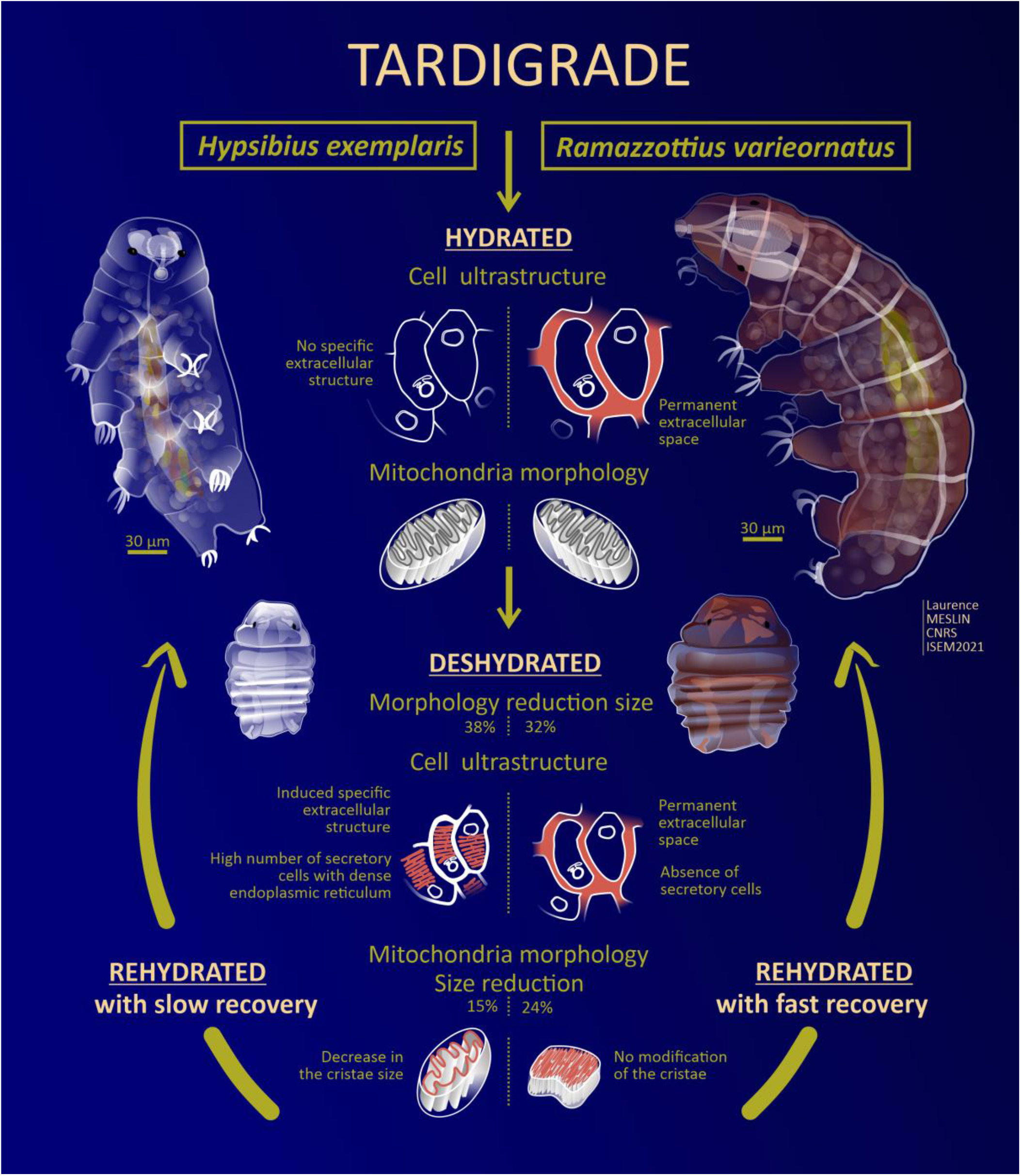
Comparative analysis of the ultrastructural divergences between *R. varieornatus* and *H. exemplaris* during the dehydration and anhydrobiote formation processes partially adapted from 57. Abstract design: Laurence Meslin CNRS, ISEM 2023.

The presence of active secretory cells during the formation of the *H. exemplaris* anhydrobiote was suggested to be associated with the production of a specific extracellular structure surrounding each cell [64]. In agreement with this observation, we were unable to detect this specific extracellular structure in the anhydrobiote (Fig. 2) of *R. varieornatus*. Moreover, es is always present in *R. varieornatus*, anhydrobiotes and hydrated specimens, and its pre-existence could explain the high capacity of *R. varieornatus* to resist anhydrobiosis stress, its speed to reach the tun stage without preconditioning and its speed of anhydrobiotic exit with an active form.

Tardigrades utilize specific IDP (TDP) gene products to survive desiccation [24,30,32]. While dehydration preconditioning has been shown to induce the upregulation of many TDPs in *H. exemplaris* and *Paramacrobiotus richtersi*, tardigrade species such as *Milnesium tardigradum* show high and constant levels of TDPs [32], which can organize to form protective noncrystalline amorphous solids inside or outside cells.

The shape of mitochondria in *R. varieornatus* and *H. exemplaris* anhydrobiote cells represents another divergent ultrastructural characteristic between the two species (Fig. 5). Compared to those in the hydrated groups, the anhydrobiotic mitochondria of *H. exemplaris* exhibited a reduced size (15%) and a decreased cristae size [64], while those of *R. varieornatus* showed a slightly greater size reduction (24%) but comparable cristae (Fig. 5a versus d). This cristae size difference between *R. varieornatus* and *H. exemplaris* may explain the respiration reactivation and faster anhydrobiotic exit of *R. varieornatus* anhydrobiotes.

In this work, we have shown that the internal ultrastructures of individual *R. varieornatus* anhydrobiotes are slightly different from those in active hydrated individuals, which contrasts with a previous report showing the neosynthesis of a specific extracellular structure associated with deep internal ultrastructural modifications in anhydrobiotic *H. exemplaris* individuals compared to hydrated individuals [64].

It is possible that the removal of the specific extracellular structure from desiccated *H. exemplaris* during the anhydrobiosis exit may slow the entire rehydration process, while *R. varieornatus*, lacking a detectable equivalent anhydrobiosis-specific ultrastructure, may not be influenced by the same way during the rehydration.

In summary, the desiccation process of *R. varieornatus* does not appear to be equivalent to that of *H exemplaris*. These differences may at least partially explain the significant differences in desiccation resistance between both species.

To date, numerous species of tardigrades have been described [1], and variations in their strategies for stress resistance, including desiccation, have been reported [68]. We have herein shown that *R. varieornatus* and *H. exemplaris*, which belong to the same Eutardigrada class, exhibit ultrastructural, morphological and dynamic divergences regarding their entry into and exit from anhydrobiosis. In addition to the comparative genetic analyses, the observations presented in this report are an additional tool to better specify the various tardigrade desiccation resistance strategies, which can be demonstrated even within the same evolutionary class.

## Acknowledgments

We thank the Montpellier MRI-RIO Imaging platform (MRI-DBS Optique and MRI-CoMET) (Montpellier, France). We thank Basile Gerbaud for his colorization of the SEM images of *Ramazzottius varieornatus*.

## Supplementary Materials

**Figure S1:**
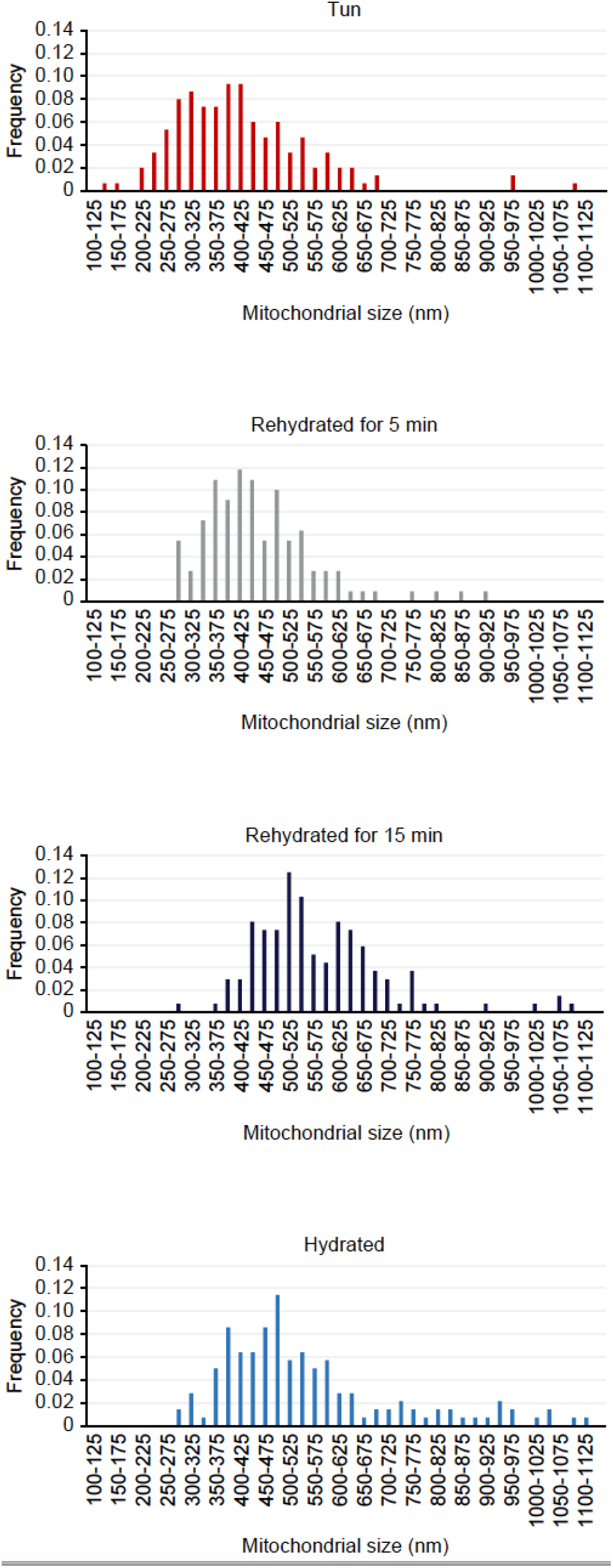
Distribution of mitochondrial size frequency depending on the tardigrade status

